# Epithelial flow into the optic cup facilitated by suppression of BMP drives eye morphogenesis

**DOI:** 10.1101/010165

**Authors:** Stephan Heermann, Lucas Schütz, Steffen Lemke, Kerstin Krieglstein, Joachim Wittbrodt

## Abstract

The transformation of the oval optic vesicle to a hemispheric bi-layered optic cup involves major morphological changes during early vertebrate eye development. According to the classical view, the lens-averted epithelium differentiates into the retinal pigmented epithelium (RPE), while the lens-facing epithelium forms the neuroretina. We find a 4.7 fold increase of the entire basal surface of the optic cup.

Although the area an individual RPC demands at its basal surface declines during optic cup formation, we find a 4.7 fold increase of the entire basal surface of the optic cup. We demonstrate that the lens-averted epithelium functions as reservoir and contributes to the growing neuroretina by epithelial flow around the distal rims of the optic cup. This flow is negatively modulated by BMP, which arrests epithelial flow. This inhibition results in persisting neuroretina in the RPE domain and ultimately in coloboma.

---

The bi-layered vertebrate optic vesicles are formed by the bilateral evagination of the late prosencephalon, a process that in teleosts is driven by single cell migration (Rembold et al., 2006). The transition of the oval optic vesicle to a hemispheric bi-layered optic cup involves major morphological transformations. In the classical view, the lens-averted epithelium of the optic vesicle differentiates into the retinal pigmented epithelium (RPE), while the lens-facing epithelium gives rise to the neuroretina, which is subsequently bending around the developing lens (Chow and Lang, 2001; Fuhrmann, 2010; Walls, 1942). This neuroepithelial bending is driven by the basal constriction of lens-facing retinal progenitor cells (RPC) (Martinez-Morales et al., 2009)(Bogdanović et al., 2012), which ultimately reduces the space an individual RPC is demanding. However, we observed that at the same time the basal optic cup surface area increases 4.7 fold (Fig. 1A-C). To identify the origin of this massive increase we performed in vivo time-lapse microscopy in zebrafish at the corresponding stages (Fig. 1D-L, movie S1) in a transgenic line expressing a membrane coupled GFP in retinal stem and progenitor cells (Rx2::GFPcaax).

**Figure 1:**
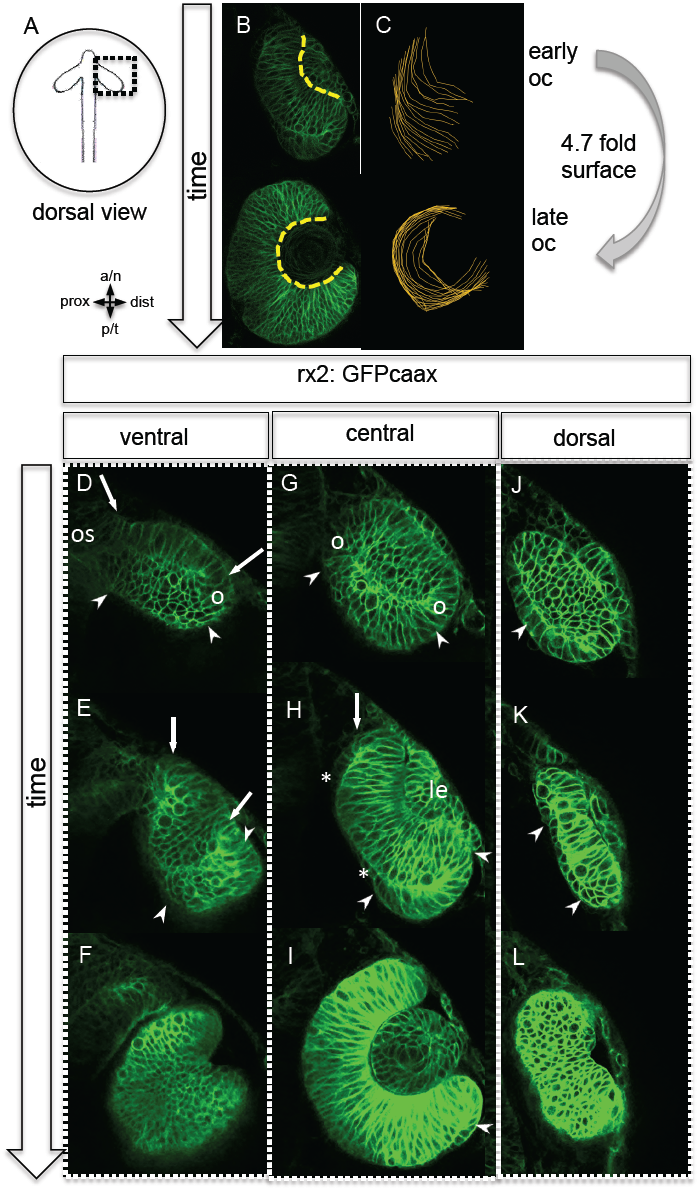
Neuroretinal surface increases during optic cup formation by epithelial flow (**A**) scheme showing the orientation of the pictures presented in B to L, (**B**) and (**C**) increase in surface from early to late optic cup stage (dashed yellow line in B, orange lines in C as overlay of optical planes), although RPCs undergo basal constriction during optic cup formation the surface increases 4.7 fold from early to late optic cup stage, (**D**) to (**L**) transition from optic vesicle to optic cup over time, shown at a ventral (D-F), a central (G-I) and a dorsal (J-L) level. The membrane localized GFP is driven by an rx2 promoter (rx2::GFPcaax), which is active in RPCs, the optic vesicle is bi-layered (D, G, J) with a prospective lens-facing (arrows in D and E) and a prospective lens-averted (arrowheads in D, G, J) epithelium, connected to the forebrain by the optic stalk (asterisk in D), at a ventral level both are connected at the distal site (circle in D), at a central level both are connected distally and proximally (circles in G), notably the morphology of the lens-averted epithelium at a dorsal level is different from central and ventral levels (arrowhead in J), over time at ventral and central levels (D-F and G-I respectively) the lens-averted epithelium is being integrated into the forming optic cup (arrowheads in D, E, G, H and I), a patch of cells in the lens-averted domain gives rise to the RPE (asterisks in H), le: developing lens, os: optic stalk

Strikingly, and in contrast to the classical view (Chow and Lang, 2001; Fuhrmann, 2010; Walls, 1942) our analysis shows that the entire bi-layered optic vesicle, gives rise to the neural retina (Fig. 1D-I), with the marked exception of a small lens-averted patch. The majority of the lens-averted epithelium (Fig. 1D and G, between arrowheads) serves as a neuro-epithelial reservoir, which is eventually fully integrated into the lens-facing neuro-epithelium (movie S1). This occurs by a sheet-like flow of lens-averted cells into the forming optic cup (Fig. 1E, H). This epithelial flow is independent of cell proliferation (Fig. S1) as demonstrated by aphidicholine treatment. It is highly reminiscent of gastrulation movements and explains the marked increase in the lens-facing basal neuroretinal surface area. Notably, a small domain of the lens-averted epithelium exhibits a different morphology and behaviour. As optic cup formation proceeds, it flattens, enlarges, shows the morphological characteristics of RPE and eventually ceases expressing RX2, a marker for retinal stem and early progenitor cells (Fig 1H, asterisks).

Our data highlight that almost the entire optic vesicle is giving rise to the neural retina. This new perspective on optic cup formation raised the question how the elongated oval optic vesicle is transformed into the hemispheric optic cup. We addressed this by following sparsely labelled cells during optic cup formation (Fig. 2A). Concomitant with lens formation we found prominent epithelial flow around the temporal and to a lesser degree nasal perimeter into the forming optic cup in line with initial previous observations (Kwan et al., 2012)(Picker et al., 2009). We uncover that the direction of the epithelial flow primarily establishes two distinct neuroretinal domains (nasal and temporal) separated by the static dorsal and ventral poles (Fig. 2D, 3A). The prospective RPE remains in the lens-averted domain and expands concomitant with the bi-furcated flow of the neuroretina from the lens-averted into the lens-facing domain (Fig. 2A, B). To address the origin of the transformation of the elongated, oval optic vesicle into the hemispheric optic cup, we quantified cellular movements along the dorsoventral axis. We found that the most prominent movements occurred in the ventral domain (Fig. 2C). Key for the formation of the ventral neuroretina is the forming optic fissure at the ventral pole of the optic vesicle. Lens-averted epithelium is flowing through this fissure into the forming optic cup to constitute the ventral neuroepithelium (Fig. 2D). Taken together we present a new model of optic cup formation, driven by gastrulation like epithelial flow from the lens-averted into the lens-facing epithelium of the forming optic cup. The epithelium flows in two domains around the temporal and nasal rim respectively and through the optic fissure of the forming optic cup. This has far reaching implications for the establishment of the retinal stem cell niche in the ciliary marginal zone (CMZ) (Centanin et al., 2011) the distal rim of the optic cup/retina.

**Figure 2:**
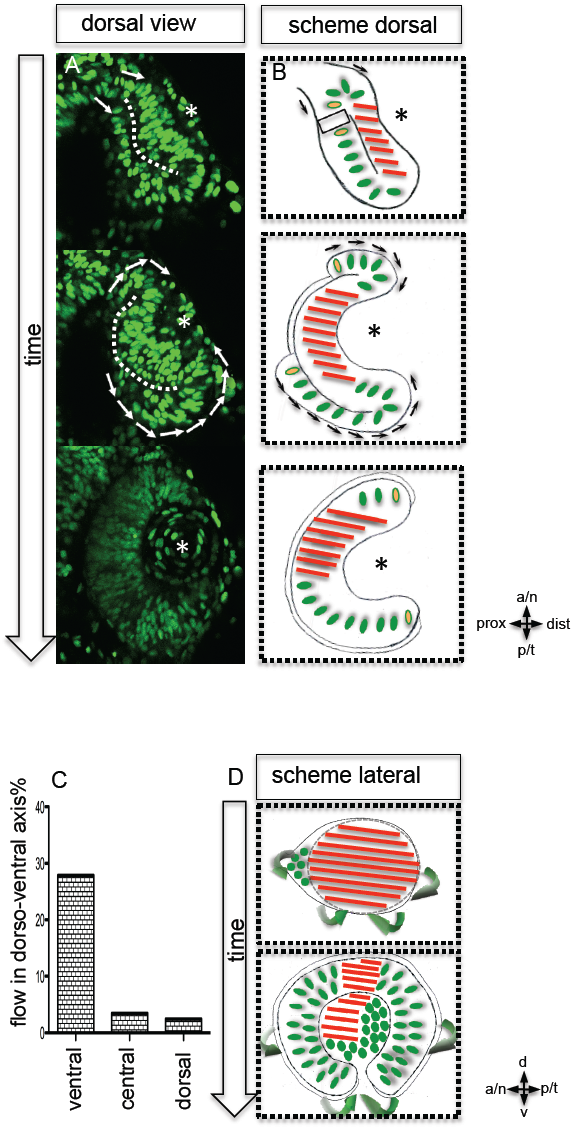
Neuroepithelial flow drives morphological changes from optic vesicle to optic cup, the role of the optic fissure and the impact on the forming stem cell domain (**A**) dorsal view on optic cup development over time visualized by nuclear GFP (H2BGFP), while the lens-facing neuroepithelium is starting to engulf the developing lens (asterisk), the lens-averted epithelium is largely integrated into the lens-facing epithelium by flowing around the distal nasal and temporal rims (arrows), a white dotted line marks the border between lens-facing and lens-averted epithelium, (**B**) scheme showing the key findings of A, the lens (asterisk) facing epithelium is marked with red bars, the lens-averted epithelium, which over time is integrated into the lens-facing epithelium is marked with green dots (except the cells at the edges are additionally marked with a yellow core), in between the last cells, which are integrated into the optic cup, the RPE will form (B), (**C**) quantification of cellular movements with respect to the dorsoventral axis, the most prominent movements in the dorsoventral axis was observed in ventral regions of the optic cup, (**D**) scheme demonstrating the optic vesicle to optic cup transition (lateral view), notably the morphological change from the elongated oval optic vesicle to the hemispheric optic cup is driven mainly by the ventral regions (arrows mark the orientation of epithelial flow) (C and D).

To address whether the domain of the presumptive CMZ arises from a mixed population of “set-aside” progenitor cells or from a predefined coherent domain, we analyzed the transition from optic vesicle to optic cup in 3D over time (4D) (movie S2). By tracking individual cells, we identified the origin of the distal retinal domain, the future CMZ, as two distinct domains (nasal and temporal) within the lens-averted epithelium at the optic vesicle stage (Fig. 3A, B, movie S3).

**Figure 3:**
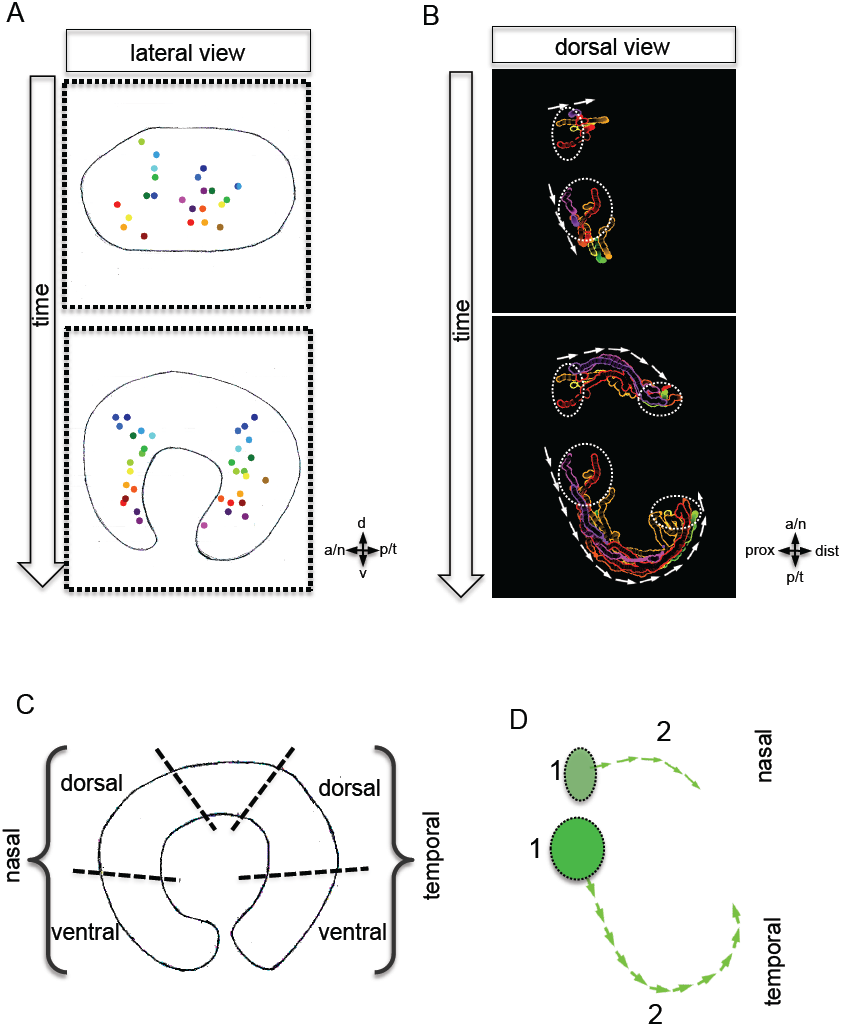
Development of the CMZ and quantification of the flow towards this domain (**A**) establishment of the presumptive CMZ domain (dorsal view), nuclear tracking of cells (maximum projection) from the lens-averted domain (encircled in upper picture) eventually residing in the forming CMZ (additionally encircled in lower picture), (**B**) scheme, lateral view over time, including the results of nuclear tracking from the presumptive CMZ back in time to the lens-averted epithelium, remarkably two distinct domains became apparent within the lens-averted epithelium as the source for the presumptive CMZ. (C) Scheme showing the optic cup from the lateral side. For quantification four distal domains were selected, nasal-dorsal, nasal-ventral, temporal-dorsal and temporal ventral. Note that the dorsal distal domain is only assembled secondarily and the ventral pole shows the optic fissure. (D) Based on differential effective distance, effective speed and directionality the migration distance was divided in three phases 1-3 in the nasal and temporal domain respectively.

To address the characteristics of the flow in its respective domains we measured the distances from the origin of the cells within the lens-averted domain towards their final destination (Fig. 3C, table) and determined their speed. It became evident that cells in the temporal, especially the temporal-ventral, domain travelled over a longer distance than the nasal domains (table). To achieve a coordinated, synchronous arrival in CMZ, the flow in the temporal domain was faster than in the nasal domain (Table 1). Based on effective speeds, distances and directionality we identified two distinct phases during the flow from the lens-averted domain towards the CMZ (Fig. 3D). In phase one the cells are highly motile, however, show only little directionality. In phase two a fast and directed flow is established that ultimately drives cells to the rim of the forming optic cup (Table 1, Fig. 3D). Notably, the onset of the fast and directed flow is seen first in the temporal and only later in the nasal domain.

As indicated above the dorsal pole of the optic vesicle remains static (Fig. 2D). Thus, the presumptive dorsal CMZ domain either originates from lens-facing neuroretina or, alternatively, is established secondarily at a later time point, like the ventral CMZ in the region of the optic fissure. The identification of lens-averted domains as the source of the future nasal and temporal CMZ is consistent with the hypothesis of a distinct origin of retinal stem cells. Our data support a scenario, in which the entire optic vesicle is initially composed of stem cells that at the lens-facing side respond to a signal to take a progenitor fate. This facilitates the tight coupling of morphogenesis with the timing of determination. Lens-averted stem cells consequently are exposed to that signal latest and are thus retaining their stem cell fate. Alternatively stemness could be maintained actively at the interface to the RPE or by a combination of both scenarios.

We demonstrated that for the neuroretinal flow to occur cell motility and thus tissue fluidity are a prerequisite. Interestingly, tissue mobility during heart jogging was recently demonstrated to be controlled by BMP antagonism (Veerkamp et al., 2013), where BMP has an “antimotogenic” effect. BMP signaling is important for various aspects of vertebrate eye development, like enhancement of RPE and inhibition of optic cup/ neuroretina development (Fuhrmann et al., 2000; Hyer et al., 2003; Müller et al., 2007; Steinfeld et al., 2013), dorso-ventral axis formation in the eye (Behesti et al., 2006; Holly et al., 2014; Koshiba-Takeuchi et al., 2000; Sasagawa et al., 2002) or the induction of the optic fissure (Morcillo et al., 2006). However, it seems crucial for specific regions of the developing eye to block BMP signaling by the expression of a BMP antagonist (Sakuta et al., 2001).

In the context of the optic vesicle to optic cup transformation we have identified follistatin a (fsta), a BMP antagonist (Thompson et al., 2005), expressed mainly in the temporal region of the optic vesicle (Fig. 4A, Fig S2), the domain with the most prominent neuroretinal flow during optic cup formation.

**Figure. 4:**
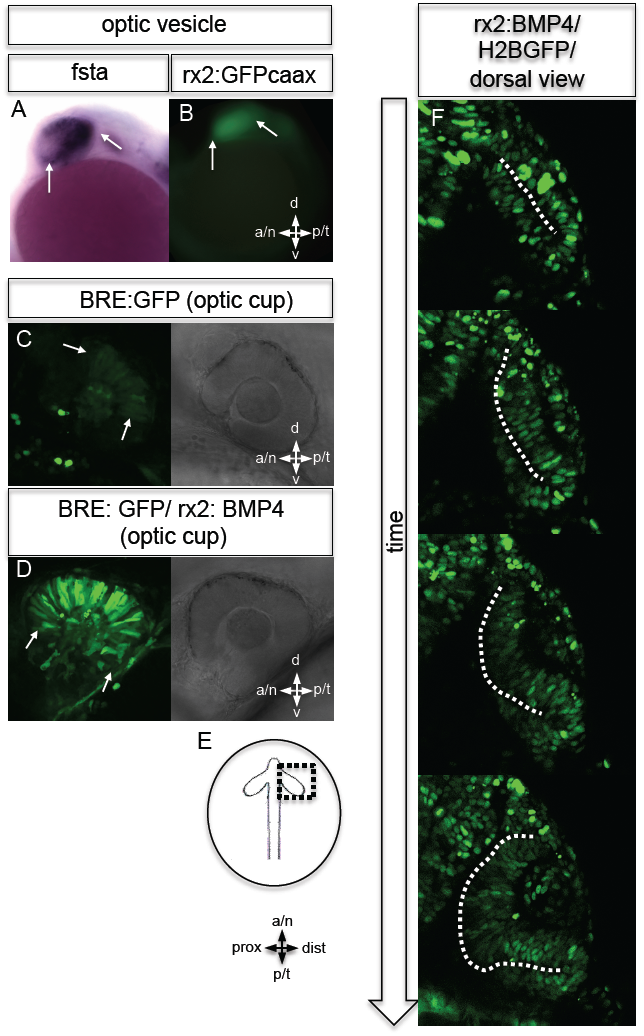
BMP antagonism drives neuroepithelial flow during optic cup formation (**A**) whole mount in-situ hybridization for fsta, please note that fsta is expressed in the optic vesicle (arrows), however, mainly in the temporal domain, (**B**) GFP expressed in the optic vesicle (arrows) of an rx2::GFPcaax zebrafish embryo, (**C**-**D**) GFP driven by the BRE and transmission/ brightfield image for orientation, strong GFP expression can be observed in the eye when BMP is driven under rx2 (arrows in D), whereas only mild GFP can be observed in controls (arrows in C), (**E**) scheme showing the orientation of the pictures presented in F, (**F**) optic cup development over time of an rx2::BMP4 embryo, cells are visualized by nuclear GFP (H2BGFP), a dotted line is indicating the border between lens-averted and lens-facing epithelium, remarkably the pan-ocular driven BMP resulted in persisting lens-averted domains.

To address the role of a graded distribution of BMP signaling we expressed BMP4 in the entire eye using an *Rx2* proximal cis regulatory element (Figs. 4B, S3A), overriding the localized BMP antagonist in the optic vesicle and optic cup. Utilizing BMP reporter fish (Laux et al., 2011) we addressed BMP signaling activity under control and experimental conditions. We did not detect BMP signaling activity early, at the optic vesicle stage (Fig S6). At the optic cup stage in contrast, moderate BMP signaling activity was observed in the dorsal retina of control fish (Fig. 4C). The pan-ocular expression of BMP4 resulted in a strong response of the reporter indicating pan-ocular BMP4 signaling (Fig. 4D).

Strikingly resembling the BMP dependent “antimotogenic” effect (Veerkamp et al., 2013), the pan-ocular BMP expression arrested epithelial flow during optic cup formation. Time-lapse *in vivo* microscopy revealed that the lens-averted part of the neuroretina persisted in the prospective RPE domain and did not contribute to the optic cup (movie S4 and S5). This ultimately resulted in an apparently ectopic domain of neuroretina that originated from a morphogenetic failure, rather than from a trans-differentiation of RPE (Fig. 5D, Fig. S3B, S4, S5). The severity of the phenotype correlated with the levels of fsta expression in the optic vesicle and was most prominent in the temporal domain of the optic vesicle. These findings highlight the importance of BMP antagonism for the epithelial fluidity during optic vesicle to optic cup transformation. We propose that the repression of BMP signaling is crucial to mobilize the lens-averted retinal epithelium to flow and eventually constitute the neural retina to a large extent.

**Figure. 5:**
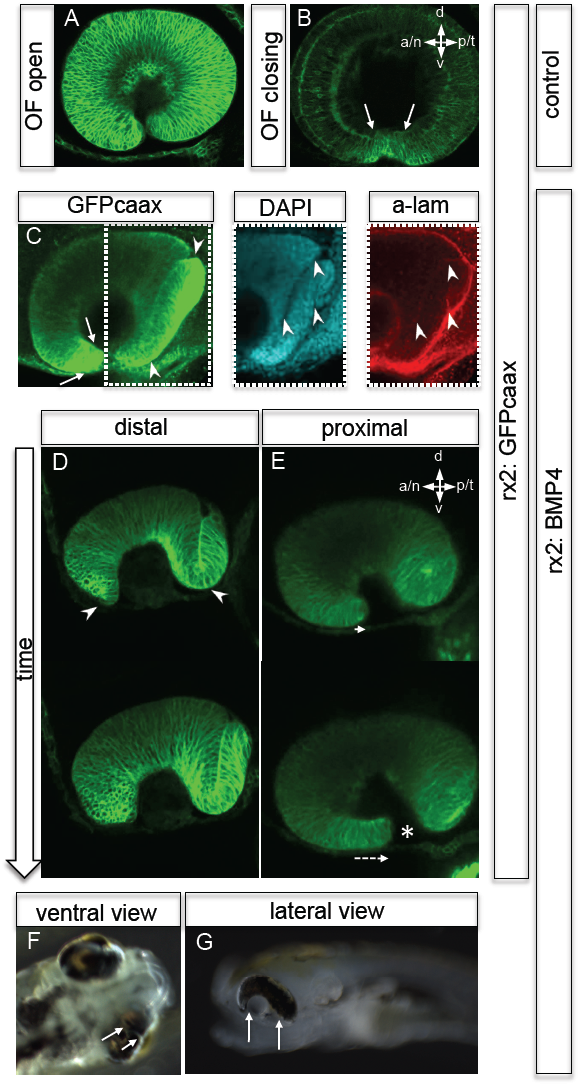
Impaired eye gastrulation results in coloboma (**A**-**B**) membrane localized GFP (rx2::GFPcaax) in a developing eye during optic fissure closure (A = early, B = late) (lateral view), rx2 is expressed in retinal stem cells/RPCs (A) and after NR differentiation becomes re-established in photoreceptors, and Müller Glia (B) while its expression is maintained in retinal stem cells of the CMZ (Reinhardt, Centanin et al, unpublished), the optic fissure margins are still undifferentiated (arrows in B), (**C**) developing eye of rx2::BMP4 fish (lateral view), membrane localized GFP (rx2::GFPcaax, anti-GFP immunointensified), DAPI nuclear labelling and anti-laminin immunostaining, the optic fissure is visible, noteworthy the temporal retina is mis-shaped and folded into the RPE domain (best visible in DAPI, arrowheads), and located on a basal membrane (arrowheads in antilaminin), especially the temporal optic fissure margin (arrowheads in GFPcaax) is located in the folded part of the temporal retina and not orderly facing the optic fissure (arrows in GFPcaax), (**D**-**E**) impaired optic fissure closure in rx2::BMP4 embryos over time at a proximal (E) and a distal (D) level, membrane localized GFP (rx2::GFPcaax), importantly next to the affected temporal optic cup also the nasal optic cup is mis-folded (arrowheads in D), remarkably, however, the nasal optic fissure margin extents into the optic fissure (dashed arrow in E) but the temporal optic fissure margin does not, likely being the result of the intense mis-bending of the temporal optic cup, this results in a remaining optic fissure (asterisk in E), (**F**-**G**) brightfield images of variable phenotype intensities observed in rx2::BMP4 hatchlings.

We further investigated the implications of the impaired epithelial flow for the successive steps of eye development (e.g. fate of the optic fissure). After initiation of neuroretinal differentiation in control embryos the undifferentiated domains are restricted to the un-fused optic fissure margins and the forming CMZ. Both can be visualized by *Rx2* (Fig. 5B, A showing Rx2::GFPcaax prior to neuroretinal differentiation). Impairment of neuroretinal flow, however, resulted in a misorganization of the optic fissure. Here, the undifferentiated *Rx2* expressing domain was found at the ultimate tip of the lens-averted neuroretinal domain that failed to flow into the optic cup and persisted in the prospective RPE (Fig. 5C). As a result of this, especially the temporal optic fissure margin was not extending orderly into the optic fissure (Figure 5D-E). This also holds true, but to a lesser extent, to the nasal optic fissure margin (Fig. 5D). Thus, the two fissure margins cannot converge resulting in a persisting optic fissure, a coloboma. Macroscopically the pan-ocular expression of BMP4 results in phenotypes with variable expressivity ranging from a “Plattauge” (Fig. 5G) in which the ventral part of the eye is strongly affected, to a milder phenotype (Fig. 5F), in which the ventral retina is developed, however, with a persisting optic fissure.

It was previously shown that an exposure of the developing eye to BMP results in a dorsalization, concomitant with a loss of ventral cell identities, likely being the cause for coloboma (Barbieri et al., 2002; Behesti et al., 2006; Koshiba-Takeuchi et al., 2000). Our data, however, conclusively show that early BMP4 exposure arrests neuroepithelial flow, resulting in a morphologically affected ventral retina while ventral cell fates remain unaltered (Fig. S7). Remarkably, the remaining lens-averted domain, which was not orderly integrated into the optic cup, eventually differentiates into neuroretina (Fig. S3B), indicated by vsx1 (Kimura et al., 2008; Shi et al., 2011; Vitorino et al., 2009) and vsx2 (formerly Chx10) expression (Vitorino et al., 2009). Notably a localization of neuroretina within the RPE domain could be mistaken for an RPE to neuroretina transdifferentiation, as proposed for other phenotypes (Araki et al., 2002; Azuma et al., 2005; Sakaguchi et al., 1997).

Taken together, our data clearly show that during optic vesicle to optic cup transformation the lens-averted part of the optic vesicle is largely integrated into the lens-facing optic cup by flowing around the distal rim of the optic cup including the forming optic fissure. Our data have far reaching implications on the generation of the retinal stem cell niche of teleosts, as the last cells flowing into the optic cup will eventually constitute the CMZ. We identify a part of the lens-averted epithelium as the primary source of the RPE. The arrest of neuroepithelial flow by the “antimotogenic” effect of BMP (Veerkamp et al., 2013) results in coloboma and thus highlights the importance of the flow through the fissure for the establishment of the ventral optic cup.

As far as the regulation of epithelial flow by fsta/BMP is concerned, the combined wt and gain of function data provide evidence for BMP to affect directionality and persistence of the epithelial flow in an inverted manner.

It does so by “freezing” motility at high BMP concentrations, facilitating strongest neuroepithelial flow in the absence of BMP signaling activity. We present a model where directionality and velocity are consequently influenced by the activity of the BMP antagonist that prevents the BMP induced “freezing” of the epithelium.

It is unlikely that the bending of the neuroretina provides the motor for the epithelial flow since it persists even in the absence of optic cup bending in the opo mutant (Bogdanović et al., 2012). Consequently, forces established outside of the neuroretina are likely to drive the flow. One tissue potentially involved is the mono-layered forming RPE. We speculate that it contributes to the flow by changing its shape from a columnar to a flat epithelium, massively enlarging its surface. This remains an interesting point, in particular given that epithelial flow is maintained even in the absence of cell proliferation in both, neuroretina and RPE.

---

**Figure S1 (related to Figure 1)**

Epithelial flow is independent of cell division

Retinal cell division was blocked by application of Aphidicolin, a well established DNA polymerase inhibitor. We addressed the epithelial flow of Aphidicolin treated wildtype embryos, injected with H2BGFP RNA at the one cell stage. Here the embryo was preincubated with Aphidicolion 5 hours prior to the start of imaging. Aphidicolin efficiently blocked cell proliferation without affecting the epithelial flow. As a low level side effect of Aphidicolin we observed cell death, in line with previous reports, importantly not affecting the epithelial flow.

**Figure S2 (related to Figure 4)**

Whole mount in situ hybridization (follistatin a (fsta)) presented as whole mount (left) and section (right)

**Figure S3: (related to Figure 4 and 5)**

(**A**) construct to drive BMP4 in RPCs, Tol2 (T2) sites flanking an rx2 promoter, driving BMP4 and a cmlc promoter driving GFP (heart) (for identification of positive embryos), (**B**) postembryonic eye development of rx2::BMP4 hatchlings, although the temporal optic cup is largely malformed and folded it can be seen clearly, that vsx1 as well as vsx2 transgenes (intensified by wholemount immunohistochemistry) are expressed in the folded epithelium (arrows) indicating at least a partial correct differentiation into neuroretinal tissue.

**Figure S4: (related to Figure 5)**

Lateral view on optic cup development over time, rx2::GFPcaax (control) is compared to rx2::BMP4 at proximal and distal levels, while in controls the lens-averted domain (yellow dotted line) is orderly integrated into the developing optic cup it persists in rx2::BMP4 embryos, note the increasing optic fissure (arrows) in rx2::BMP4 embryos.

**Figure S5: (related to Figure 5)**

Dorsal view on optic cup development of an rx2::BMP4 embryo over time at ventral versus central/ dorsal levels, the yellow dotted line indicates the border between the lens-facing and the lens-averted domain, remarkably the lens-averted domain is not orderly integrated into the optic cup (compare to figure 1 and 2), notably an altered morphology of the ventral optic vesicle can be observed showing the domain which is not going to be integrated orderly (arrows).

**Figure S6:** No active BMP signaling during optic vesicle formation

This data was obtained by analyzing BMP signaling reporter fish at the optic vesicle stage. Analysis of signaling activity shows that BMP signaling is absent from the optic vesicles at this stage of development, visualized by the in vivo reporter. Note signaling activity in other tissues such as the tail-bud.

**Figure S7:** Ventral retinal cell identity unaltered in response to rx2::BMP4

Whole mount in situ hybridization shows that vax2 remains expressed in the forming ventral optic cup even if BMP4 is expressed panocularly (rx2::BMP4), clearly showing that morphogenesis is not affected due to an early fate switch into “all dorsal”.

**Table**

Table presenting quantified data for cellular movements during optic vesicle to optic cup transition. Phase 2 (see Figure 3D) marks the onset of a smooth flow movement.

**movie S1: (related to Figure 1)** (control) optic vesicle to optic cup transition visualized with rx2::GFPcaax (orientation as in Fig. 1)

**movie S2: (related to Figure 2)** (control) optic vesicle to optic cup transition visualized by H2BGFP RNA into rx2::GFPcaax (orientation as in Fig. 2)

**movie S3: (related to Figure 2)** (control) optic vesicle to optic cup transition visualized by H2BGFP RNA into rx2::GFPcaax (orientation as in Fig. 2), data as in movie S2 with tracked cells (maximum projection) to the presumptive CMZ

**movie S4: (related to Figure 4)** (rx2::BMP4) optic vesicle to optic cup transition visualized by H2BGFP RNA (orientation as in Fig. 3)

**movie S5: (related to Figure 4)** (rx2::BMP4) optic vesicle to optic cup transition visualized by rx2::GFPcaax (orientation as in Fig. 3)

## Experimental Procedures

### Transgenic zebrafish

BMP4 was cloned via directional Gateway from zebrafish cDNA into a pEntr D-TOPO (Invitrogen) vector with the following primers: forw: 5’ CACCGTCTAGGGATCCCTTGTTCTTTTTGCAGCCGCCACCATGATTCCTGGTAATCGAATGCTG 3’, rev: 5’ TTAGCGGCA GCCACACCCCTCGACCAC 3’.

The expression construct was assembled via a Gateway reaction using Tol2 destination vector containing a cmlc: GFP, a 5’Entry vector containing an rx2 promoter, the vector containing the BMP4 and a 3’Entry vector containing a pA sequence. The construct was injected into zebrafish eggs.

The rx2::GFPcaax construct was assembled via Gateway (Invitrogen) and injected into zebrafish eggs. The BRE: GFP zebrafish line was kindly provided by Beth Roman. The Vsx1: GFP zebrafish line was kindly provided by Lucia Poggi. The Vsx2: RFP zebrafish line was kindly provided by the lab of William Harris.

## Quantification of optic cup surface

Optic cup surfaces were measured with the help of FIJI (Image J NIH software). The mean of the length of the measured lines (Fig. 1C) of two adjacent optical sections was multiplied by the optical section interval.

## Microscopy

Confocal data of whole mount immunohistochemical stainings a Leica SPE microscope was used. Samples were mounted in glass bottom dishes (MaTek). Olympus stereomircoscope was used for recording brightfiled images of rx2::BMP4 hatchlings and the overview of the expression of rx2::GFPcaax. For whole mount insitu data acquisition a Zeiss microscope was used. Time-lapse imaging was performed with a Leica SP5 setup which was upgraded to a multi photon microscope (Mai Tai laser). It was recorded in single photon modus and multi photon modus. For timelapse imaging embryos were embedded in 1% low melting agarose and covered with zebrafish medium, including tricain for anesthesia. Left and right eyes were used and oriented to fit the standard dorsal view or view from the side.

## Whole mount in situ hybridization

Whole mount in situ hybridization for fsta AB/AB zebrafish at ov stage were used. The procedure was largely performed according to Quiring et al (Quiring et al., 2004).

## Whole mount immunohistochemistry

Immunohistochemistry was performed according to a standard whole mount immunohistochemistry protocol. Briefly, embryos/ hatchlings were fixed, washed, bleached (KOH/H_2_O_2_ in PTW), and blocked (BSA, DMSO, TritonX100, NGS, PBS). Samples were incubated in primary antibody solution (anti-laminin, Abcam) (anti-GFP, life technologies) (anti-dsRED, Clontech ) in blocking solution. Samples were washed and incubated in secondary antibody solution with DAPI added. Consecutively samples were washed and mounted for microscopy.

### Quantitative Analyses

Quantification of dorso-ventral movement:

The amount of movement in the dorso-ventral axis was quantified using a supervoxel based Optical Flow algorithm (Amat et al., 2013). The pixel wise output was visualized by applying a spherical coordinate system to the eye using a custom made ImageJ plugin. The colour coding is based on the sign of the polar angle theta and the sign of the azimuth angle phi, as well as on their respective combinations. The quantification was performed by counting the labelled pixels in an ImageJ macro.

### Cell tracking

Cells were tracked manually using MtrackJ (Meijering et al., 2012) in Fiji (ImageJ) (Schindelin et al., 2012) back in 4D stacks to their original location or until lost. Only tracks with a significant length were used for the visualizations. Centered on the track cells are represented as spheres. Partially results are presented in a side view where the dorso-ventral axis originally represented as the z-axis has now become the y-axis. A factor of 10.5703 is introduced in order to adjust the data of the former z-axis to the other two axes. The colour coding is done by choosing colours from an 8 bit Look up table and applying them from the dorsal to the ventral side based on the end of the track. Partially tracking results are presented as tailed spheres. The spheres are based on the tracking data using an average over the last three timepoints. The image is stretched in the z-axis using a factor of 10.5703, to adjust the scale to the x and y axes. Tails are created using a lookup table with 16 different shades per colour. The respective shade is defined by the distance and difference in time between the recent position and the position on the tail.

## Acknowledgements

We want to thank Lea Schertel for excellent technical assistance, members of the Wittbrodt lab for material (P. Haas and A. Schmidt for the rx2: GFP construct and zebrafish line, B. Höckendorf and M.Stemmer for H2BGFP mRNA, B. Höckendorf for FIJI related input) and constructive stimulating feedback, B. Roman for the BRE: GFP zebrafish, William Harris for the vsx2: RFP zebrafish line and Lucia Poggi for making the vsx1: GFP zebrafish line available. LS is recipient of a fellowship from the Hartmut-Hoffman-Berling International Graduate School (HBIGS). This work was supported by the Deutsche Forschungsgemeinschaft (JW, KK, SL) and the ERC (JW).

